# Multi-scale mapping along the auditory hierarchy using high-resolution functional UltraSound in the awake ferret

**DOI:** 10.1101/249417

**Authors:** Célian Bimbard, Charlie Demené, Constantin Girard, Susanne Radtke-Schuller, Shihab Shamma, Mickael Tanter, Yves Boubenec

**Affiliations:** Audition Team, Laboratoire des Systèmes Perceptifs CNRS UMR 8248, École Normale Supérieure, PSL Research University, Paris, France; Institut Langevin, CNRS UMR 7587, Inserm U979, ESPCI ParisTech, PSL Research University, Paris, France; University of Maryland College Park, MD, USA; equal contribution

## Abstract

A major challenge in neuroscience is to longitudinally monitor whole brain activity across multiple spatial scales in the same animal. Functional UltraSound (fUS) is an emerging technology that offers images of cerebral blood volume over large brain portions. Here we show for the first time its capability to resolve the functional organization of sensory systems at multiple scales in awake animals, both *within* structures by precisely mapping sensory responses, and *between* structures by elucidating the connectivity scheme of top-down projections. We demonstrate that fUS provides stable (over days), yet rapid, highly-resolved 3D tonotopic maps in the auditory pathway of awake ferrets, with unprecedented sharp functional resolution (100μm). This was performed in four different brain regions, including small (1-2mm^3^ size), subcortical (8mm deep) and previously undescribed structures in the ferret. Furthermore, we used fUS to map longdistance projections from frontal cortex, a key source of sensory response modulation, to auditory cortex.

## Introduction

Functional ultrasound imaging (fUS) based on Ultrafast Doppler was first introduced in neuroimaging in 2011 ^1^. Using ultrasonic plane wave emissions, this system exhibits a 50-fold enhanced sensitivity to blood volume changes compared to conventional ultrasound Doppler techniques ^2^. Relative to fMRI, it has much faster temporal dynamics (ms rather than seconds), substantially higher spatial resolution (100μm rather than millimeters), lower cost, and greater portability. However, most fUS studies thus far have investigated its sensitivity in capturing coarse-grained sensory responses ^3–5^, or used it to explore indirect inplane brain connectivity ^6,7^. Here, we pushed the limits of fUS imaging to demonstrate its capability in capturing a *fine-grained* functional characterization of sensory systems and *direct, long-distance* connectivity scheme between brain structures. We show that fUS imaging can rapidly produce highly-resolved 3D maps of responses reflecting precise tonotopic organizations of the vascular system in the almost complete auditory pathway of awake ferrets. We further demonstrate that fUS imaging can provide voxel to voxel independent information (100μm;), indicative of its extreme sensitivity and spatial functional resolution. These measurements are conducted over several days in small (1-2mm^3^ size) and deep nuclei (8mm below the cortical surface), as well as across various fields of the auditory cortex. On a broader scale, we describe how fUS can be used to assess long distance (out-of-plane) connectivity, with a study of top-down projections from frontal cortex to the auditory cortex. Therefore, fUS can provide a multi-scale functional mapping of a sensory system, from the functional properties of highly-resolved single voxels, to inter-area functional connectivity patterns.

## Results and discussion

Physiological experiments were conducted in 3 awake ferrets *(Mustela putorius furo*, thereafter called V, B and S). After performing craniotomies over the temporal lobe, chronic imaging chambers were installed (both hemispheres in one animal, and right hemispheres in the other two) to access a large portion of both the auditory (middle and posterior ectosylvian gyri - resp. MEG and PEG) and visual cortex (in caudal suprasylvian and lateral gyri) (Fig. 1a). The 3D scan of the craniotomy via Ultrafast Doppler Tomography ^10^ revealed the indepth vasculature of the Auditory Cortex (AC) surrounded by the supra-sylvian sulcus (Fig. 1a and b). In addition, we were able to detect and image deep auditory-responsive structures such as the Medial Geniculate Body (MGB), the Inferior Colliculus (IC) and the Lateral Lemniscus (LL), as well as light-responsive nuclei such as the Lateral Geniculate (LGN) (Figure 1—figure supplement 1).

In order to reveal the tonotopic organization of the auditory structures, we recorded in each voxel the evoked hemodynamic responses to pure tones of 5 different frequencies, and then computed the resultant 3-dimensional tonotopic map (Fig. 1c-e, Figure 1—figure supplement 2). Within a relatively short time (10 to 15 minutes per slice), we could accurately reproduce the known tonotopic organization of the primary (A1, MEG) and secondary auditory cortex (PEG), with a high‐ to low-frequency gradient in A1, reversing to a low‐ to high-frequency gradient in the dorsal PEG (Fig. 1c). We note that the fUS enabled us to map within the challenging deep folds of the ferret auditory cortex, such as the suprasylvian sulcus (sss) and pseudo-sylvian sulcus (pss). Recordings could be performed in the same slice across days, with a high repositioning precision (error <1 slice, 200μm in that case), which was within the range of the out-of-plane point-spread function for fUS (Figure 1—figure supplement 3).

Large-scale, 3D functional maps were also recorded in the deep and smaller structures of the auditory thalamus (MGB, Fig. 1d), the inferior colliculus (IC, Fig. 1e) and the lateral lemniscus (LL, Fig. 1e). The 3D views obtained in fUS allowed us to describe for the first time the tonotopical organizations of the ferret’s MGB and LL. This is particularly remarkable in the latter structure in which we characterize a precise tonotopic map despite its small size (~1mm-long) and subcortical position (8mm deep below brain surface). Moreover, such a large field of view allows one to measure simultaneously the functional organization of any coplanar structure (such as A1 and the MGB here), thus opening the door to precise, frequency specific (thalamo-cortical) connectivity studies. In this respect, future development of high frequency fUS matrix-probes for 3D UfD imaging ^11^ will extend this capability to any brain structure.

**Figure 1.**
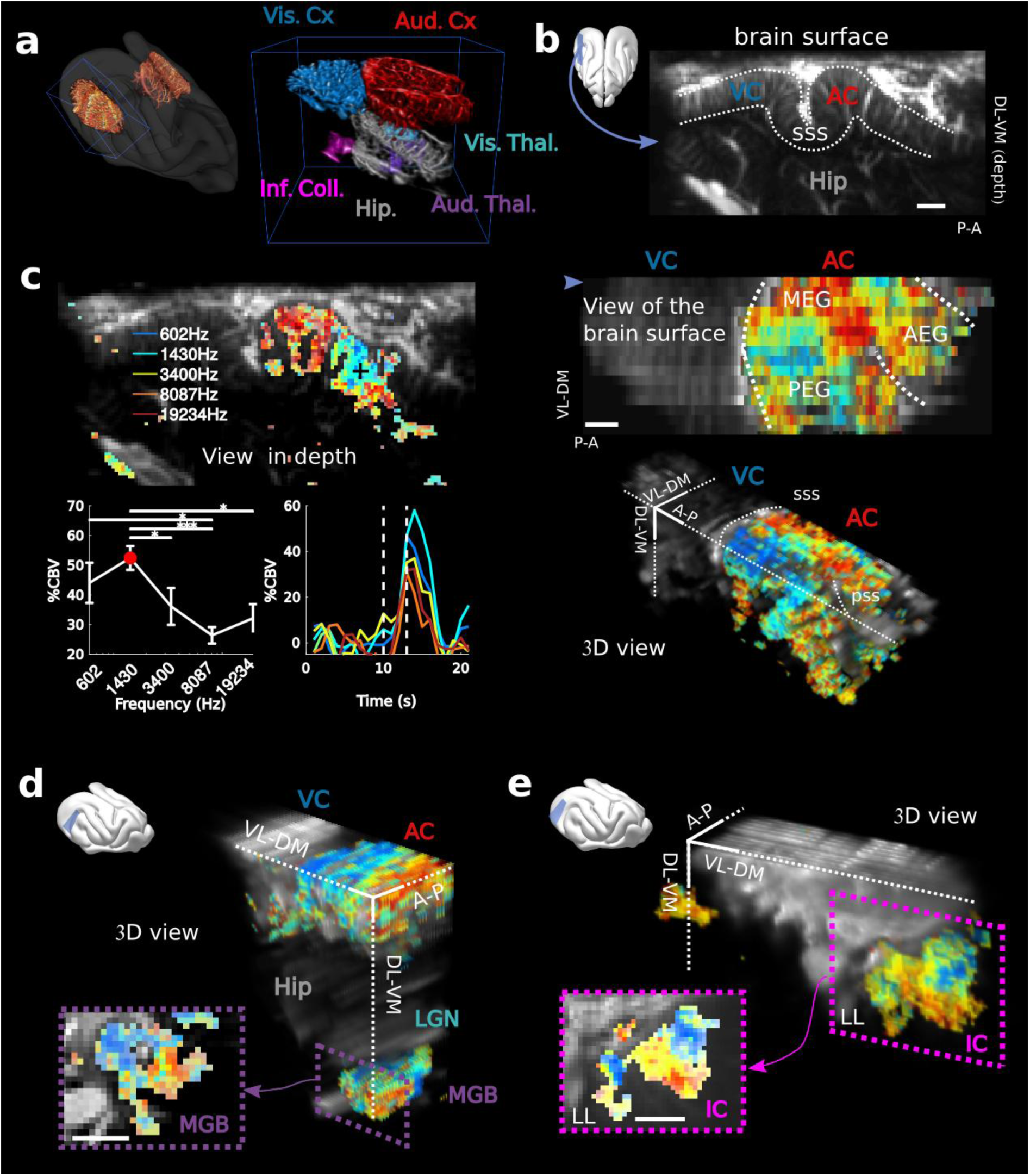
fUS imaging reveals the tonotopic organization of cortical, sub-cortical, and intracortical auditory structures in the awake ferret. (a). Left: UFD-T of the left and right craniotomies, superimposed on an fMRI scan of a ferret brain. Right: magnification of the blue bounding box (left). Auditory structures: auditory cortices (AC), medial geniculate body (MGB), inferior colliculus (IC). Other structures: hippocampus (Hip), visual cortex (VC). (b). Structural view of a tilted parasagittal slice (~30° from D-V axis) of the visual and auditory cortices (represented as a blue plane on the 3D brain). Lining delineates the cortex. (c) Upper left: Tonotopic organization of the slice described in (b). Lower left: tuning curve (mean±sem) and average responses in %CBV (see Online Methods) for the voxel located in the upper panel (black cross). Upper right: combination of 16 similar slices over the surface of the AC, arrow depicts slice of (b). Lower right: 3D reconstruction of the whole AC’s functional organization. (d) 3D reconstruction of both the auditory cortex and auditory thalamus (non-tonotopic areas were masked on this reconstruction for clarity of the representation). Inset: single slice centered on the MGB. Its tonotopic axis runs along the PL-AM axis. Note that (b-d) were extracted from the left side of the brain, but flipped for visual clarity and coherence. (e) 3D reconstruction of the inferior colliculus and the lateral lemniscus (LL). Inset: single slice centered on the IC. Both (d) and (e) are tilted coronal slices (~30° from D-V axis). Their tonotopic axis runs along a ~20°-tilted D-V axis All individual and converging scale bars: 1mm. D: dorsal, V:ventral, M: medial, L: lateral, A: anterior, P: posterior Fig 1 ‒ figure supplement 1: **Responses to visual and auditory stimuli in the cortex and thalamus.** Fig 1 ‒ figure supplement 2: **Tonotopies in AC, IC and MGB for other animals**. Fig 1 ‒ figure supplement 3: **fUS allows for high recording stability and repositioning over days.**

Single-trial analysis is essential for understanding brain dynamics and behavioral variability. However, it remains a challenge as it necessitates to record high-quality signal from a large number of neurons/voxels at the same time. In order to estimate the reliability and selectivity of fUS single-trial responses, we used MultiVoxel Pattern Analysis (MVPA) to decode the stimulus frequency from the hemodynamic signal. Using a simple linear decoder, we attained high decoding accuracy in the auditory cortex (from 0.46 to 0.63 probability, with chance at 0.2) which was even more striking in the IC and LL (from 0.72 to 0.98), despite their smaller size and subcortical location (Fig. 2a). These results suggest that single trials show reliable and significant activity across all structures.

On a different scale, we sought to demonstrate whether fUS could also reveal encoding differences across cortical layers. We focused on imaging the small vessels in the cortex (keeping only data corresponding to an axial projection of blood flow lower than 3.1 mm/s) and defined cortical layers using an unfolding algorithm providing a flattened version of the AC (Figure 2—figure supplement 1). A linear decoder yielded a significantly higher decoding accuracy when using only measurements at intermediate cortical depths (p<1e-3), peaking around 400-500μm below the surface (up to 0.83, mean 0.67), consistent with it being granular. As a control, we note that baseline blood volume and response magnitude did not show a similar depth-dependent profile (Figure 2—figure supplement 1), suggesting that the observed decoding accuracy could reflect primarily functional differences in the neuronal encoding of frequency across cortical layers ^12^ or variations in capillaries structure within cortical layers ^13^. Importantly, all these results could be confirmed in single slice recordings, and over several days (Figure 2—figure supplement 2), showing that the hemodynamic signal imaged in fUS is reliable enough to decode brain activity on a single-trial basis within a single experiment.

Next, we took a closer look at the tonotopic organization in different structures to examine how tuning curves in neighboring voxels change abruptly. This finding exemplifies the ability of fUS imaging to measure independent information at a very small spatial scale. To quantify the minimal functional spatial resolution of the technique, we defined a discriminability index between voxels, and focused on sharp transition areas (Fig. 2b left panels). We found that fUS can discriminate responsiveness of neighboring voxels, with a functional resolution as fine as 100μm (Fig. 2b). Furthermore, we were able to discriminate voxels based on their tuning curves within a distance of 300μm in as little as 20 repetitions per frequency (Fig. 2b). These results suggest that fUS can be useful to assess the fine organization of vascular domains within brain structures and to better understand the functional coupling between local neuronal activity and the dynamics of surrounding blood vessels, two important questions for hemodynamic-based techniques ^9,14^.

**Figure 2:**
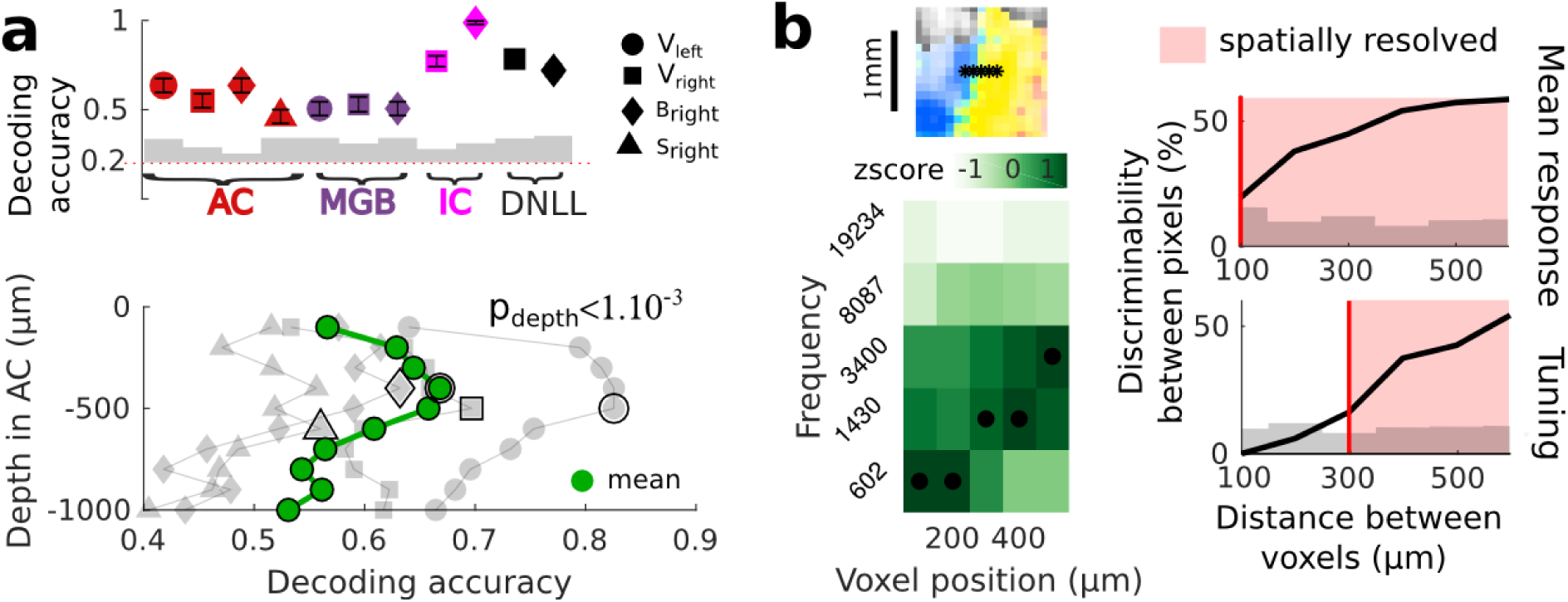
Key features of fUS in awake animals: decoding accuracy, layer effect, and effective spatial resolution. (a) Upper panel: Decoding accuracy over the 5 frequencies, in different structures and different craniotomies (see legend). Grey histogram shows the upper limit for chance (p<1e-2, mean±2sem computed over 10 randomized decoding sessions). All structures showed significant decoding (p<1e-5). Lower panel: decoding accuracy over depths, computed from the activity in the AC of 4 different animals (grey plots). All showed a similar profile, with the accuracy peaking between 400 and 500μm. The green plot shows the average trend (repeated-measure ANOVA over depth, p<1e-3). (b) Upper left: zoom on a sharp tonotopic transition from low to medium frequency. Black stars represent consecutive voxels. Lower left: heatmap of the z-scored tuning curves of the consecutive voxels, with the best frequency indicated by a black dot, showing a shift from low to medium frequency preference. Right: quantification of the lower spatial limit at which one can significantly find significant differences in the responsiveness (upper) or tuning (lower) of two voxels, with respect to their distance. Grey histogram shows the upper limit for chance (p<5e-2, 5% percentile over 50 randomizations). In that specific case, it was respectively 100μm and 300μm. Fig 2 ‒ figure supplement 1: **Controls for the decoding across depths.** Fig 2 ‒ figure supplement 2: **Single-slice recordings show high decoding possibility on an actual single-trial basis.**

Another fundamental view of brain function and functional organization is revealed by mapping brain connectivity among various structures. Localizing and quantifying such connections in awake animals, however, remains technically challenging since tracer injections are not an option, and fMRI gives only access to indirect, spatially diffuse measures of connectivity strength. Here, we demonstrate that fUS can be used to probe the functional connectivity between two brain structures that are far apart: the frontal and the auditory cortices. The frontal cortex (FC) is a region that has been shown to be involved in top-down modulation of early sensory areas, and in particular of the auditory cortex ^15,16^. To reveal its potential links to the auditory areas, we electrically stimulated at different points within the FC while recording evoked hemodynamic responses in the auditory cortex of an awake (slightly sedated) animal (Fig. 3a). Importantly, this technique does not require any precise priors on the location and nature of the terminal projections. By imaging widely in the auditory cortex, we observed evoked activity in the fundus of the PSS (PSSC/insula), which was maximal for a certain depth and position of the stimulating electrode (Fig. 3b, Figure 3— figure supplement 1). By contrast, there was no evoked activity recorded in secondary auditory areas such as the PEG (Figure 3—figure supplement 2). We also observed a suppression of blood flow in the MEG, possibly originating from polysynaptic connections between FC and A1 ^17^. Anatomical confirmation of the existence of such descending projections from FC to PSSC/insula was independently obtained from anterograde tracer injections in FC, which revealed monosynaptic projections that targeted the PSSC/insula (Fig. 3c), consistent with the functional connectivity pattern revealed by the fUS approach. Because the neighboring regions have been reported to be multimodal ^18,19^, we subsequently explored the responsiveness of the FC-targeted PSSC/insula to acoustic and visual stimuli. We found this region to be poorly responsive to broadband noise, and not driven by visual stimuli (Figure 3—figure supplement 3). Altogether, this experiment offers a proof-of-concept of how fUS can serve as a tool to characterize large-scale functional connectivity without sacrificing any resolution. For example, one may explore subtle connectivity changes during the course of learning, possibly combined with optogenetic tools to target specific neuronal subpopulations.

**Figure 3:**
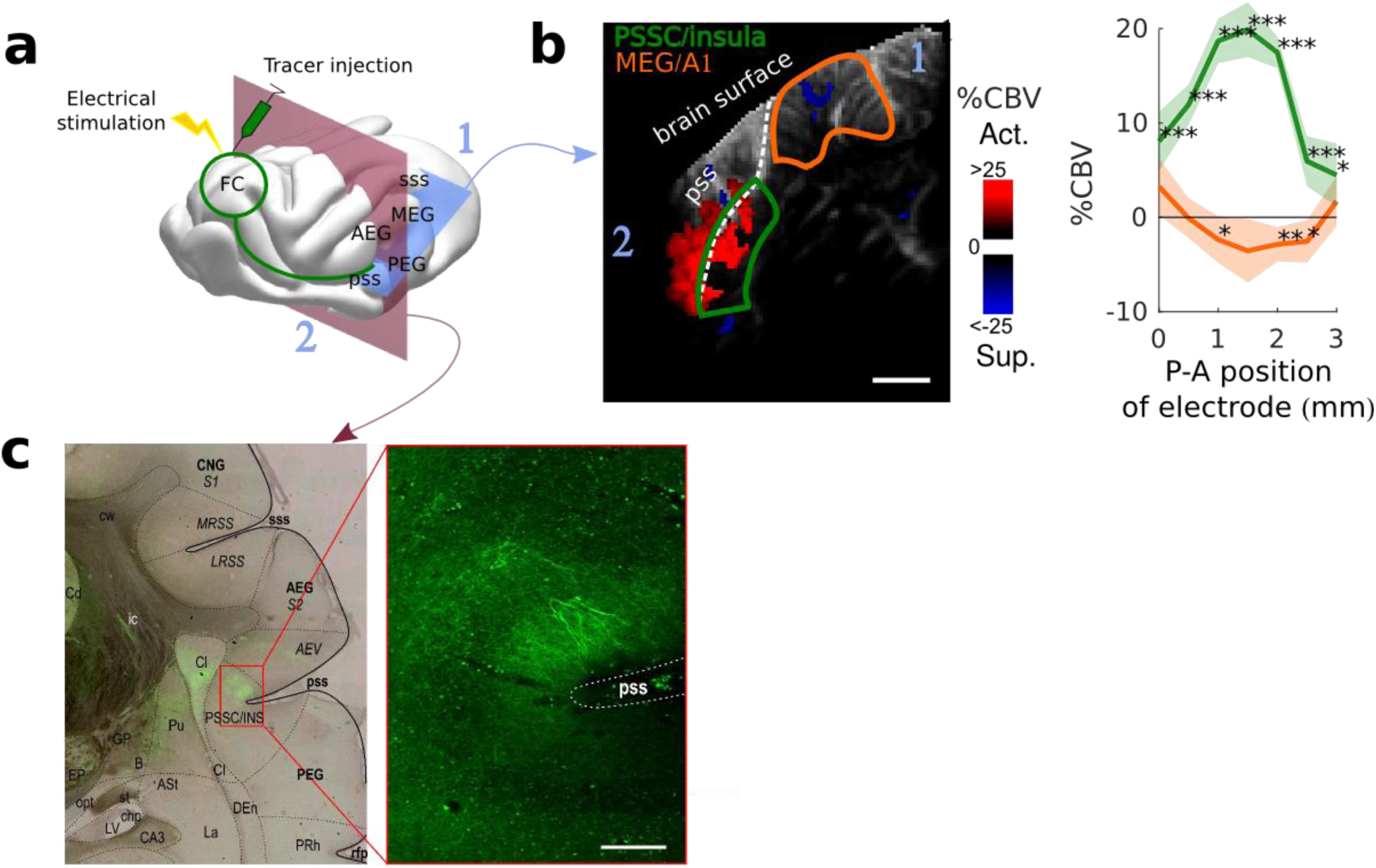
Exploring long-distance connectivity: the example of top-down projections from dlFC to the auditory system. (a) Ferret brain with localization of electric stimulation (lightning) and virus (tracer) injection site (green circle) with symbolized projections, site of fUS imaging shown in (b) (blue plane) and coronal slice represented in (c) (red plane). (b) FC-AC direct projection patterns revealed in fUS. Left: fUS imaging plane along the PSSC/insula, showing modulations of hemodynamic activity in MEG (orange delimitation) and PSSC/Insula (green delimitation) evoked by FC stimulation (map thresholded at +4sem). The numbers 1 and 2 are here to help orientation. Right: %CBV in the 2 regions of interest after FC electric stimulation (highlighted in the left panel) with respect to the postero-anterior position of the stimulation electrode (0 represents 25.5mm from caudal crest, 3 represents 28.5mm), revealing a hot-spot of connectivity at about 1mm (i.e 26.5mm from caudal crest) (mean±2sem). (c) Anatomical confirmation of connectivity. Left: bright field combined with fluorescence imaging, showing green fluorescent FC projections concentrated in the depth of the PSSC/insula and delineated anatomical structures (scale: 200μm). Right: close-up of the labelled FC projection terminals in the fundus of the pss (PSSC/insula). Fig 3 ‒ figure supplement 1: **Frontal Cortex - Auditory cortex connectivity explored further: cortical depth**. Fig 3 ‒ figure supplement 2: **Frontal Cortex - Auditory cortex connectivity explored further: secondary areas**. Fig 3 ‒ figure supplement 3: **Frontal Cortex - Auditory cortex connectivity explored further: sound and vision**.

To conclude, we have shown that fUS imaging can serve as a technique to record in awake animals an exceptionally stable (over days), high-resolution and simultaneous mapping of various brain regions, be they large, small, superficial, or deep. This was done over multiple scales, from functional tuning of individual voxels to large-scale connectivity between brain regions. The mapping is both rapid and precise, as illustrated by the ease with which single-trial information can be decoded from its high-sensitivity signal. Furthermore, fUS can be a valuable tool in acquiring broad, yet accurate views of the functional organization of unmapped brain regions and their connectivity with the rest of the brain. Finally, fUS imaging can be readily adapted to mobile and highly stable configurations ^20^, which will make it ideally suited for behavioral cognitive neuroscience studies requiring extended observations, as in the characterization of the neural correlates of learning.

## Acknowledgements

We thank Marc Gesnik from the Institut Langevin for his valuable inputs for the ultrasound sequence programming and Roberto Toro for the ferret fMRI scan. This work was supported by ANR-10-LABX-0087 IEC et ANR-10-IDEX-0001-02 PSL* and research grants from the European Research Council under the European Union’s Seventh Framework Program (FP7/2007-2013) / ERC Advanced grant agreement n° 339244-FUSIMAGINE and ERC Advanced grant agreement n° ADG_20110406-ADAM and R01-DC005779 (SS). The project received the technical support of the INSERM Technology Research Accelerator in Biomedical Ultrasound.

## Online methods

### A. Animal preparation

Experiments were approved by the French Ministry of Agriculture (protocol authorization: 01236.02) and strictly comply with the European directives on the protection of animals used for scientific purposes (2010/63/EU). To secure stability during imaging, a stainless steel headpost was surgically implanted on the skull and stereotaxis locations of the dorsolateral frontal cortex (FC) and the auditory cortex (AC) were marked ^1^. Under anaesthesia (isoflurane 1%), 4 craniotomies above the auditory cortex were performed on 3 ferrets, using a surgical micro drill, yielding a ~15x10mm window over the brain. After clean-up and antibiotic application, the hole was sealed with an ultrasound-transparent TPX cover, embedded in an implant of dental cement ^2^. Animals could then recover for one week, with unrestricted access to food, water and environmental enrichment.

For fUS imaging, animals were habituated to stay in a head-fixed contention tube. The ultrasonic probe was then inserted in the implant and acoustic coupling was assured via degassed ultrasound gel. Experiments were conducted in a double-walled sound attenuation chamber. All sounds were synthesized using a 100 kHz sampling rate, and presented through Sennheiser IE800 earphones (HDVA 600 amplifier) that was equalized to achieve a flat gain. Stimulus presentation were controlled by custom software written in Matlab (MathWorks).

### B. Ultrafast Doppler imaging

We used a custom miniaturized probe (15 MHz central frequency, 70% bandwidth, 0.110 mm pitch, 128 elements) inserted in a 4 degree-of-freedom motorized setup. The probe was driven using a custom fully-programmable ultrasonic research platform (PI electronics) and dedicated Matlab software.

#### 1) 3D vascular imaging

Vascular anatomy of the brain portion accessible from the craniotomy was imaged in 3D using the Ultrafast Doppler Tomography (UFD-T) strategy described in ^3^. Briefly, this method acquires 2D Ultrafast Power Doppler (UfD) images at a frame rate of 500 Hz. Each frame is a compound frame built with 11 tilted plane wave emissions (-10° to 10° with 2° steps) fired at a PRF of 5500Hz, combined with mechanical translation and rotation, and then postprocessed via a Wiener deconvolution to correct for the intrinsic out-of-plane loss of resolution, so that we ultimately recover an isotropic 100μm 3D resolution. In the end, a 3D (14x14x20mm) blood volume reconstruction of the vasculature is obtained (voxel size: 50μm, isotropic resolution 100μm). This 3D vascular imaging was performed on each craniotomy, and was used as a local reference framework, specific to the craniotomy, where recording planes could be repositioned over days using correlation methods.

#### 2) fUS imaging

fUS imaging relies on rapid acquisition (every 1s) of ultrasensitive 2D Power UfD images of the ferret brain. For each Power image, 300 frames are acquired at a 500 Hz frame rate (covering 600ms, i.e. one to two ferret cardiac cycles), each frame being a compound frame acquired via 11 tilted plane wave emissions (-10° to 10° with 2° steps) fired at a PRF of 5500 Hz. Image reconstruction is performed using an in-house GPU-parallelized delay-and-sum beamforming. Those 300 frames are filtered to discard global tissue motion from the signal using a dedicated spatio-temporal clutter filter^4^ based on a singular value decomposition of the spatio-temporal raw data. Blood signal energy (called Power UfD) is then computed for each voxel (100x100x~400μm, the latter dimension, called elevation, being slightly dependent of depth) by taking the integral 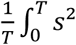 over the 300 time points ^5^ This power Doppler is known to be proportional to blood volume ^6^. A certain band of Doppler frequencies can be chosen before computation of the power using a bandpass filter (in our case a 5^th^ order low-pass Butterworth filter), enabling the selection of a particular range of blood flow speeds, i.e. discriminating between capillaries and arterioles (slow blood flow) and big vessels (fast blood flow). In our study, we set the filtering to better focus on small vessels with axial velocity lower than 3.1mm.s^-1^ when indicated in the text.

### C. Protocol for sensory response acquisition

Auditory responses were studied by playing different sounds through animal earphones during recording of the brain activity via fUS imaging. The protocol for sound presentation is as follows: 10 seconds of silence (baseline), then 2 seconds of sound followed by 8 seconds of silence (return to baseline). Visual responses were obtained by playing a flickering red-light stimulus instead of sound, with the same durations of different epochs.

1) Localization of the auditory structures: In order to find the boundaries of the auditory structures in the imaged portion of the brain, white noise sound was played (70dB).
2) Mapping of the tonotopic organization of the auditory structures; Auditory structures are known to exhibit tonotopic organization based on extensive physiological and structural studies (in the ferret, see ^7–11^). To image these tonotopic maps, we played unmodulated pure tones while recording fUS images at five equally spaced frequencies on a logarithmic scale (602 Hz, 1430 Hz, 3400 Hz, 8087 Hz, 19234 Hz, covering the auditory hearing spectrum of the ferret, at 65 dBSPL). The tones were played in random order, 10 trials/frequency (20 in the animal S.). To obtain the whole tonotopic organization in a 3D volume, this process was repeated in different slices in order to build a 3D stack from successive 2D slices (spaced by 300μm). Each slice was acquired in ~15 minutes, thus allowing us to map in 3D the whole auditory cortex within a few hours. We note that these tone stimuli elicited large and reliable responses in the whole auditory tract despite being unmodulated. This suggests that a variety of other auditory stimuli (such as natural sounds) can be used to elicit stronger responses and hence reveal more organizational properties.

### D. FC stimulation

FC electric stimulations were adapted from previously described protocols ^12,13^. Platinium-iridium stimulation electrodes (impedance 200-400kOhms, FHC) were positioned in the FC using stereotaxic coordinates, obtained from functional recordings in behaving animals (AP: 25.5-28.5mm (0 to 3 mm on Fig. 2d) from caudal crest / ML: 2mm ^14^). Each trial consisted of 10s of baseline, then 6s of monophasic stimulation at 100Hz and 200μA (2ms pulses, 200ms-long train, repeated at 2Hz), after a return to baseline of 10s. The %CBV was computed as the mean response between 3 and 6s after stimulation onset. 30 trials were performed for each A-P position of the electrodes. In these connectivity experiments, the animal was slightly sedated using a small dose of medetomidine (Domitor 0.02mL at 0.08 mg.kg^-1^) to reduce movement artifacts.

### E. Anatomical tracers

A one year old female ferret weighing 620g received a 2μl injection of pAAV2.5-CaMKIIa-hChR2(H134R)-EYFP (PennCore) as anterograde tracer into left FC. Six months later the animal was perfused and the brain was cryoprotected, shock frozen and cut on a cryostat into 50 μm thick frontal sections into parallel series of which one was counterstained with neutral red. For overview images, combined brightfield and fluorescence images were taken with a Hamamatsu slide scanner 2.0HT (Institut de la Vision) (Fig. 2e, left). For details, fluorescence images were taken with a virtual slide microscope (VS120 S1, Olympus BX61VST) at 10* magnification (Fig. 2e, right). Anatomical structures were reconstructed in accord with the ferret brain atlas ^14^.

### F. Signal processing, analysis& statistics

#### 1) %CBV

Power UfD signal was normalized towards the baseline to monitor changes in Cerebral Blood Volume (%CBV). This quantity varied after stimulus presentation (Fig. 1c) and we quantified voxel responses with the mean of %CBV in a time-window 3 to 5s after sound onset. Tonotopy of the imaged structures was mapped as follows: for each voxel this mean vascular response across the 5 tested frequencies was used to determine its best frequency (BF). Statistical differences of the responses to different frequencies in an individual voxel (Fig. 1c, tuning curve) were assessed using a Wilcoxon rank sum test (post-hoc test after significant ANOVA p<1e-3). Maps were thresholded by showing only voxels that had (i) a minimal 15% response and (ii) a mean response at their BF highly correlated (p<1e-3) with the mean hemodynamic response. This mean hemodynamic response was used to approximate the typical vascular response to stimulus (as the Hemodynamic Response Function does for fMRI) and was computed in each structure as the average response over all the voxels showing a response to sound with z-score>3. Note that thresholds could be adjusted depending on the overall responsiveness of different structures and different animals, for illustration purpose. Last, maps were spatially smoothed with a 3x3x1 voxel gaussian filter (std = 0.5). The view of the brain surface (Fig. 1c) was computed as the mean BF averaged from 5 to 10 voxels from the auditory cortex surface delimited manually. For 3D reconstructions of the cortex only, manually adjusted masks were used in order to show only tonotopic regions, and avoid crowdy representations caused by voxel transparency in the 3D visualization. Cortical depths were obtained by manually tracing the surface (just below the pia’s blood vessels) and depth limits of the cortex. The 10 different depths were then automatically extracted by a custom-made algorithm (Fig. 2a and Figure 2—figure supplement 1 & 2). The number of voxels at each depth was then equalized for the decoding analysis.

For the single slice analysis presented in. Figure 2—figure supplement 2, the protocol was designed to speed up tone-responses acquisition (2s tone, and random interval of 4 to 6s - uniformly distributed - between two tone-presentations). We then used a General Linear Model (GLM) to compute impulse responses of individual voxels to each tone frequency, without any predefined hemodynamic response function. This allowed us to present more stimuli (75 per frequency) in a relatively shorter time (~45 min).

#### 2) Decoding

Frequency selectivity of the auditory cortex was assessed using a 5-class linear classifier and a leave-one out strategy: for each frequency pair, vascular responses of the 2 frequencies (%CBV averaged over 4 to 5s after sound onset) were separated in a voxel-based space via a linear boundary optimized on 9 of the 10 trials in a learning set. Overall, pseudo-populations were built by grouping, across all slices recorded within the same structure, trials with identical frequency labels. The decoder was run over 100 shuffles of these pseudo-populations, where train and test sets were randomly chosen. In single slice analysis (Figure 2—figure supplement 2), we used a Fisher decoder (normalized by covariance) in order to take into account the noise correlation between voxels in decoding analysis. This was doable thanks to the higher number of tone presentations that allowed us to have a stable estimation of the covariance matrix.

In order to prove the significance of the obtained accuracy, we shuffled the responses of each voxel over all trials, and performed the same analysis. We computed the mean and variance of the random distributions with 10 randomizations, and performed an unpaired Wilcoxon rank sum test to check whether the real accuracy was significantly out of the random distribution.

To evaluate whether cortical depth had an effect on decoding accuracy (Fig. 2a), we performed a one-way repeated-measure ANOVA over the 4 different craniotomies, with depth as the factor.

#### 3) Resolution quantification

In order to quantify the minimal spatial scale at which fUS can provide independent information from two neighbouring voxels, we focused on sharp edges of functional transition and performed 2-way (voxel and frequency as factors) ANOVA on the tuning curves of each pair of voxels along a transect (example transect shown in Fig. 2b, top left). The voxel factor quantified the dissimilarity in the average responses for two voxels, being thus representative of an overall responsiveness dissimilarity when significant. The interaction term (frequency x voxel) quantified how dissimilar the tuning curves were for two different voxels, independently of their overall responsiveness. This term therefore represented our ability to discriminate between different functional voxel tuning. Importantly, these values depend on the smoothness of the underlying functional neuronal map (the sharper the better) and on the number of trials used in each experiments (the higher the better). Here, we show that using only 20 trials per frequency, we could go down to a functional resolution comparable to the voxel size (100μm) for the overall responsiveness, and of 300μm for the tuning.

We randomized the responses over all voxels and all frequencies and performed the same analysis to find the average distribution expected by chance for both responsiveness and tuning dissimilarity percentages. We determined the spatial resolution as the shortest distance at which the actual number of dissimilar pairs was above the 95^th^ percentile of the randomized distribution.

## References

1. Mace E., E. et al. Functional ultrasound imaging of the brain. Nat Methods 8, 662–664 (2011).

2. Mace, E. et al. Functional ultrasound imaging of the brain: theory and basic principles. IEEE Trans. Ultrason. Ferroelec., Freq. Contr. 60, 492–506 (2013).

3. Tiran, E. et al. Transcranial Functional Ultrasound Imaging in Freely Moving Awake Mice and Anesthetized Young Rats without Contrast Agent. Ultrasound Med. Biol. 43, 1679–1689 (2017).

4. Osmanski, B. F. et al. Functional ultrasound imaging reveals different odor-evoked patterns of vascular activity in the main olfactory bulb and the anterior piriform cortex. Neuroimage 95, 176–184 (2014).

5. Gesnik, M. et al. 3D functional ultrasound imaging of the cerebral visual system in rodents. Neuroimage 149, 267–274 (2017).

6. Osmanski, B.-F., Pezet, S., Ricobaraza, A., Lenkei, Z. & Tanter, M. Functional ultrasound imaging of intrinsic connectivity in the living rat brain with high spatiotemporal resolution. Nat Commun 5, 5023 (2014).

7. Rideau Batista Novais, A. et al. Transcriptomic regulations in oligodendroglial and microglial cells related to brain damage following fetal growth restriction. Glia 64, 2306–2320 (2016).

8. Logothetis, N. K. & Wandell, B. A. Interpreting the BOLD Signal. Annu. Rev. Physiol. 66, 735–769 (2004).

9. O’Herron, P. et al. Neural correlates of single-vessel haemodynamic responses in vivo. Nature 534, 378–382 (2016).

10. Demené, C. et al. 4D microvascular imaging based on ultrafast Doppler tomography. Neuroimage 127, 472–483 (2016).

11. Provost, J. et al. 3-D ultrafast doppler imaging applied to the noninvasive mapping of blood vessels in Vivo. IEEE Trans. Ultrason. Ferroelectr. Freq. Control 62, 1467–1472 (2015).

12. Guo, W. et al. Robustness of Cortical Topography across Fields, Laminae, Anesthetic States, and Neurophysiological Signal Types. J. Neurosci. 32, 9159–9172 (2012).

13. Adams, D. L., Piserchia, V., Economides, J. R. & Horton, J. C. Vascular supply of the cerebral cortex is specialized for cell layers but not columns. Cereb. Cortex 25, 3673–3681 (2015).

14. Harrison, R. V, Harel, N., Panesar, J. & Mount, R. J. Blood capillary distribution correlates with hemodynamic-based functional imaging in cerebral cortex. Cereb. Cortex 12, 225–233 (2002).

15. Fritz, J., Shamma, S., Elhilali, M. & Klein, D. Rapid task-related plasticity of spectrotemporal receptive fields in primary auditory cortex. Nat. Neurosci. 6, 1216–1223 (2003).

16. Winkowski, D. E., Bandyopadhyay, S., Shamma, S. A. & Kanold, P. O. Frontal cortex activation causes rapid plasticity of auditory cortical processing. J Neurosci 33, 18134–18148 (2013).

17. Logothetis, N. K. et al. The effects of electrical microstimulation on cortical signal propagation. Nat. Neurosci. 13, 1283–1291 (2010).

18. Bizley, J. K., Nodal, F. R., Bajo, V. M., Nelken, I. & King, A. J. Physiological and anatomical evidence for multisensory interactions in auditory cortex. Cereb Cortex 17, 2172–2189 (2007).

19. Bizley, J. K. & King, A. J. Visual-auditory spatial processing in auditory cortical neurons. Brain Res. 1242, 24–36 (2008).

20. Sieu, L.-A. et al. EEG and functional ultrasound imaging in mobile rats. Nat Methods 12, 831–834 (2015).

## References

1. Atiani, S. et al. Emergent selectivity for task-relevant stimuli in higher-order auditory cortex. Neuron 82, 486–499 (2014).

2. Sieu, L.-A. et al. EEG and functional ultrasound imaging in mobile rats. Nat Methods 12, 831–834 (2015).

3. Demené, C. et al. 4D microvascular imaging based on ultrafast Doppler tomography. Neuroimage 127, 472–483 (2016).

4. Demené, C. et al. Spatiotemporal Clutter Filtering of Ultrafast Ultrasound Data Highly Increases Doppler and fUltrasound Sensitivity. IEEE Trans. Med. Imaging 34, 2271–2285 (2015).

5. Mace, E. et al. Functional ultrasound imaging of the brain: theory and basic principles. IEEE Trans. Ultrason. Ferroelec., Freq. Contr. 60, 492–506 (2013).

6. Rubin, J. M., Bude, R. O., Carson, P. L., Bree, R. L. & Adler, R. S. Power Doppler US: a potentially useful alternative to mean frequency-based color Doppler US. Radiology 190, 853–856 (1994).

7. Moore, D. R., Semple, M. N. & Addison, P. D. Some acoustic properties of neurones in the ferret inferior colliculus. Brain Res. 269, 69–82 (1983).

8. Pallas, S. L., Roe, A. W. & Sur, M. Visual projections induced into the auditory pathway of ferrets. I. Novel inputs to primary auditory cortex (AI) from the LP/pulvinar comples and the topography of the MGN-AI projection. J. Comp. Neurol. 298, 50–68 (1990).

9. Versnel, H., Mossop, J. E., Mrsic-Flogel, T. D., Ahmed, B. & Moore, D. R. Optical imaging of intrinsic signals in ferret auditory cortex: responses to narrowband sound stimuli. J. Neurophysiol. 88, 1545–1558 (2002).

10. Nelken, I. et al. Large-scale organization of ferret auditory cortex revealed using continuous acquisition of intrinsic optical signals. J Neurophysiol 92, 2574–2588 (2004).

11. Bizley, J. K., Nodal, F. R., Nelken, I. & King, A. J. Functional organization of ferret auditory cortex. Cereb Cortex 15, 1637–1653 (2005).

12. Tolias, A. S. et al. Mapping cortical activity elicited with electrical microstimulation using fMRI in the macaque. Neuron 48, 901–911 (2005).

13. Logothetis, N. K. et al. The effects of electrical microstimulation on cortical signal propagation. Nat. Publ. Gr. 13, 1283–1291 (2010).

14. Radtke-Schuller, S. A Cyto‐ and Myeloarchitectural Brain Atlas of the Ferret (Mustela putorius) in MRI-aided Stereotaxic Coordinates. (Springer Nature, 2018).

